# The components of an electrical synapse as revealed by expansion microscopy of a single synaptic contact

**DOI:** 10.1101/2023.07.25.550347

**Authors:** Sandra P. Cárdenas-García, Sundas Ijaz, Alberto E. Pereda

## Abstract

Most nervous systems combine both transmitter-mediated and direct cell-cell communication, known as ‘chemical’ and ‘electrical’ synapses, respectively. Chemical synapses can be identified by their multiple structural components. Electrical synapses are, on the other hand, generally defined by the presence of a ‘gap junction’ (a cluster of intercellular channels) between two neuronal processes. However, while gap junctions provide the communicating mechanism, it is unknown whether electrical transmission requires the contribution of additional cellular structures. We investigated this question at identifiable single synaptic contacts on the zebrafish Mauthner cells, at which gap junctions coexist with specializations for neurotransmitter release and where the contact defines the anatomical limits of a synapse. Expansion microscopy of these contacts revealed a detailed map of the incidence and spatial distribution of proteins pertaining to various synaptic structures. Multiple gap junctions of variable size were identified by the presence of their molecular components. Remarkably, most of the synaptic contact’s surface was occupied by interleaving gap junctions and components of adherens junctions, suggesting a close functional association between these two structures. In contrast, glutamate receptors were confined to small peripheral portions of the contact, indicating that most of the synaptic area works as an electrical synapse. Thus, our results revealed the overarching organization of an electrical synapse that operates with not one, but multiple gap junctions, in close association with structural and signaling molecules known to be components of AJs. The relationship between these intercellular structures will aid in establishing the boundaries of electrical synapses found throughout animal connectomes and provide insight into the structural organization and functional diversity of electrical synapses.

## INTRODUCTION

Synapses are specialized cell-cell contacts at which relevant functional information is exchanged between two neurons, either via a chemical messenger (‘chemical synapses’) or direct cell-cell channels (‘electrical synapses’) (1). While the molecular complexity of chemical synapses with structurally distinct pre- and postsynaptic components has long been recognized (2, 3), less is known regarding the molecular and structural complexity of electrical synapses. Electrical synapses are a modality of neuronal communication mediated by structures known as ‘gap junctions’ (GJs) (4). These structures contain intercellular channels formed by the apposition of two hemichannels, each provided by one of the connected cells, and which cluster together into GJ ‘plaques’ (4). Hemichannels are formed by proteins called ‘connexins’ (4, 5) in vertebrates and ‘innexins’ (6, 7) in invertebrates that, though unrelated in sequence, share a similar membrane topology which allows them to assemble into intercellular channels. While GJs are ubiquitous and present in virtually every tissue of the organism providing metabolic coupling (4), they additionally serve as a pathway of low resistance for the spread of electrical currents between neurons (and cells of the heart), the main form of signaling in the brain, which is fast enough to operate within the time frame required for decision making by neural circuits (8–10). Electrical synapses are generally perceived as structurally simpler than chemical synapses and exclusively involving the function of intercellular channels. However, recent data indicates that the function of these channels is under the control of their supporting molecular scaffold (11–14), suggesting that neuronal GJs are complex molecular structures whose function requires the contribution of multiple molecular components (15). Such molecular complexity is likely to underlie plastic changes in the strength of electrical synapses (16, 17), which are capable of dynamically reconfiguring neural circuits (17, 18).

Thus far, investigations of the properties of vertebrate electrical transmission have solely focused on the functional properties of the channel-forming proteins, the connexins, the molecules that regulate them, and their interactions at the GJ plaque. However, could a single neuronal GJ per se be considered an electrical synapse? Alternatively, does electrical transmission rely on additional structural components? The identification of the structural components of a chemical synapse is facilitated by the presynaptic bouton, which anatomically defines its limit (19). In contrast, neuronal GJs are typically found connecting cell somata or other neuronal processes (9, 19, 20), such as dendrites and axons, making it more difficult to define the exact anatomical boundaries that constitute an electrical synapse. Interestingly, neuronal GJs can also occur at synaptic boutons, generally coexisting with specializations for chemical transmission (19, 21). This is the case for auditory afferents terminating as single ‘Large Myelinated Club Endings’ (22, 23) or ‘Club endings’ (CEs) on the lateral dendrite of the teleost Mauthner cells (a pair of large reticulospinal neurons involved in tail-flip escape responses in fish) (24, 25), each containing GJs and specializations for chemical transmission (26). Because of their experimental accessibly and functional properties, these terminals are considered a valuable model to study vertebrate electrical transmission, as they more easily allow for the correlation between structure and function of synaptic features (16, 26–28). Since the bouton marks the anatomical limits of a synapse, these contacts offer the opportunity to examine the anatomical structures that together make an electrical synapse.

Here, we used expansion microscopy (29) to expose the presence and spatial arrangement of synaptic components in CEs of larval zebrafish. CEs from larval zebrafish share comparable morphological and functional properties with those of adult fish (12, 30–32), and due to their genetic accessibility, allow the opportunity to investigate the functional link between the structures enabling electrical transmission and its regulation. Expansion revealed the presence of multiple well-defined puncta distributed throughout the contact area, which, consistent with the notion that they represent GJ plaques, each exhibited labeling for the fish homologs of the widespread mammalian connexin 36 (Cx36), Cx35 and Cx34, as well as for the GJ scaffolding protein Zonula Occludens (ZO1) (11, 12, 31, 33). Strikingly, expansion following staining with N-cadherin and ß-catenin antibodies, protein components of AJs, showed that these proteins are also distributed all throughout the contact area, but in a fashion that is mutually exclusive with connexin. This suggests that the subsynaptic topography of electrical synapses is carefully and intimately coordinated. Finally, double-labeling with Cx and glutamate receptor antibodies showed that, while GJs are distributed throughout the entire contact area, a much smaller number of glutamatergic sites are restricted to the periphery occupying a small fraction of the contact’s surface. Our data suggest that synaptic communication at electrical synapses results from not one, but the coordinated action of multiple GJs of variable size, which may require the functional contribution of additional structures, such as AJs.

## RESULTS

A group of auditory afferents, each of which terminate as a single synaptic contact, known as a Club ending (CE), on the distal portion of the lateral dendrite of the Mauthner (M-) cell (22, 23) (Fig. 1A). Because of their unusual large size and experimental accessibility, CEs represent a valued model for the correlation of synaptic structure and function. Ultrastructural analysis of CEs in adult goldfish (26, 34, 35) and larval zebrafish (30) revealed the presence of gap junctions (GJs) coexisting with specializations for neurotransmitter release. Consistent with these synaptic specializations, stimulation of CEs evokes a synaptic response that combines electrical and chemical transmission (27, 30, 32, 36). A wealth of evidence indicates that GJs at these terminals consist of heterotypic intercellular channels created by the apposition of a presynaptic hemichannel formed by connexin Cx35.5 and a postsynaptic hemichannel formed by Cx34.1, two of the Cx35 and Cx34 orthologs (31, 33). These junctions also contain the scaffolding protein ZO1 (11, 12), which regulates channel function. The synaptic areas of CEs can be visualized by immunolabeling for these GJ proteins, which are revealed as large fluorescent oval areas at the distal portion of the lateral dendrite of the M-cell (Fig.1 B-D).

**Figure 1.**
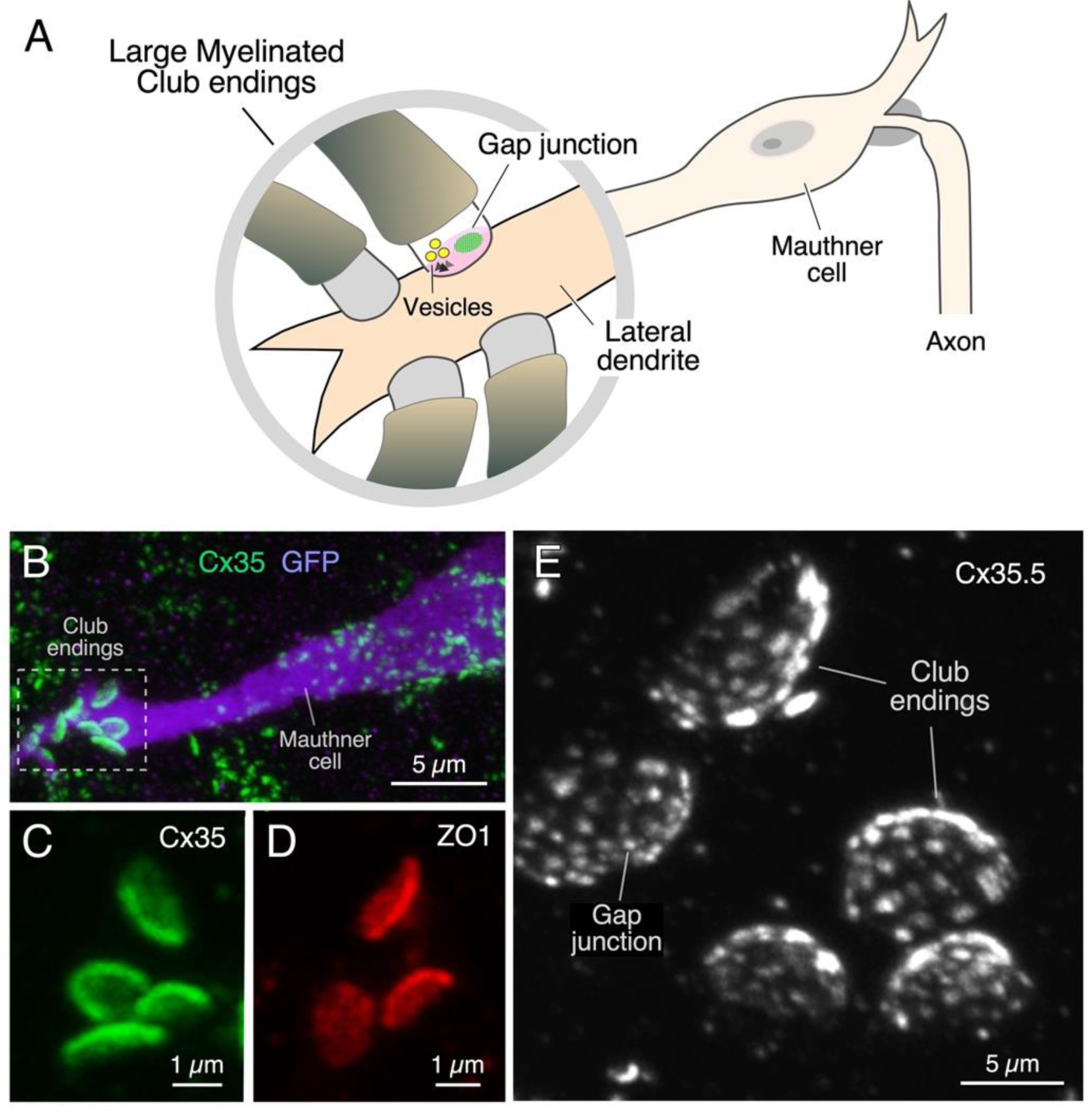
Expansion microscopy of Club ending contact areas in larval zebrafish. (A) The cartoon illustrates the spatial distribution of auditory afferents that terminate as single Club endings (CEs), each containing both gap junctions (GJs, green) and specializations for chemical transmission (vesicles), on the distal portion of the lateral dendrite of the Mauthner (M-) cell. (B) Confocal image with anti-GFP (purple) and anti-Cx35/36 (green) showing a long stretch of the lateral dendrite (projection of 34 confocal Z-sections, totaling 13.5 µm), revealing the contact areas of several CEs. (C-D) Contact areas of individual CEs labeled with anti-Cx35/36 (C, green; projection of 12 sections, totaling 4.7 µm) and anti-ZO1 (D, red; projection of 4 sections, totaling 1.6 µm). (E) Expansion microscopy (ProExM) with anti-Cx35.5 increases the size of CE synaptic contact areas, enabling the visualization of intrasynaptic components (projection of 19 sections, totaling 10.5µm).

Immunolabeling is commonly used to define the biochemical composition of a synapse by allowing detection of the presence of specific proteins. We combined this approach with a protein-retention expansion protocol (proExM, (29)) to explore, not only the presence, but the relative distribution of the structures formed by various synaptic proteins throughout the contact areas of CEs in 5 days post fertilization (dpf) zebrafish. Tissue expansion with anti-Cx35.5, revealed the presence of multiple Cx35.5-positive puncta at these contacts (Fig. 1E), reminiscent of that observed at goldfish CEs and known to correspond to GJ plaques (11, 37). The main axis of the oval synaptic contact area defined by Cx35 or ZO1 labeling increased 4-fold, from 2.11 ± 0.005 µm (n=38) to 8.25 ± 0.035 µm (n=40) in the expanded samples (mean ±SEM). Moreover, expansion led to a 13-fold increase in the area of the contact, from 2.63 ± 0.01 µm^2^ (n=40) to 35.12 ± 0.19 µm^2^ (n=46). A concern when using proExM is whether this procedure leads to the distortion of normal anatomical features. Therefore, to determine the degree of isotropy of the expansion, we measured the ratio between the short and long diameter of the CE oval areas (S/L ratio) in pre- and post-expanded samples. The short and long diameters in non-expanded tissue averaged 1.61 ± 0.004 µm (n=38) and 2.11 ± 0.005 µm (n=38), respectively, and 5.16 ± 0.022 µm (n=40) and 8.25 ± 0.035 µm (n=40), respectively, in expanded terminals. Analysis showed that the average S/L ratio decreased in the expanded tissue, with the decrease reaching statistical significance [non-expanded, 0.77 ± 0.002 (n=38); expanded, 0.64 ± 0.003 (n=40); p<0.01]. However, the decrease of the average S/L ratio was less than 18%. Moreover, the distribution of S/L ratios from expanded tissue was nearly identical to the distribution from non-expanded tissue, and both were well described by normal distributions (Supp. Fig 1). This result indicates that the expansion procedure had no selective effects on CEs, but rather affected all uniformly. The difference between expanded and non-expanded S/L ratios might also result from an underestimation of the small diameter due to a minor tilting along the long diameter in ‘En face’ views of expanded CEs, which are more difficult to obtain because of their larger size. Together with our synaptic alignment findings (see below), these results show that the expansion procedure had minimal effects on synaptic structure. Thus, expansion of these single synapses resulted in a more than 10-fold increase of the synaptic contact area, allowing for a more detailed visualization of the relative distribution of its synaptic components.

CEs combine electrical with chemical transmission. To investigate the area occupied by each form of transmission we labeled for Cx35.5 as a marker of GJs, and glutamate receptor 2 (GluR2) as a marker of glutamatergic transmission (Fig. 2A-C). As expected, colocalization analysis (Mander’s coefficient analysis) revealed that labeling for Cx35.5 and GluR2 were mutually exclusive (Fig 2D). Yet, while Cx35.5 labeling covered the majority of the area, labeling for GluR2 was substantially lower and limited to the contact’s peripheral margin (Fig. 2A-C). To quantify this differential distribution, we defined two regions of interest (ROI) at ‘En face’ views of the CE contact: one representing the central ¾ of the area (‘center’) and another representing the peripheral remaining ¼ of the area (‘periphery’). Estimates of the relative intensity of Cx35.5 vs GluR2 in each ROI showed that while GluR2 is constrained to the periphery, Cx35.5 is homogenously distributed through the contact area (Fig 2E-F). This distribution is consistent with previous EM reconstruction of CEs in adult goldfish, at which specializations for transmitter release were found to be restricted to the periphery of the contact. Thus, while GJs occupy most of the surface, chemical transmission occupies a smaller and peripheral portion, suggesting that most of the contact operates as an electrical synapse.

**Figure 2.**
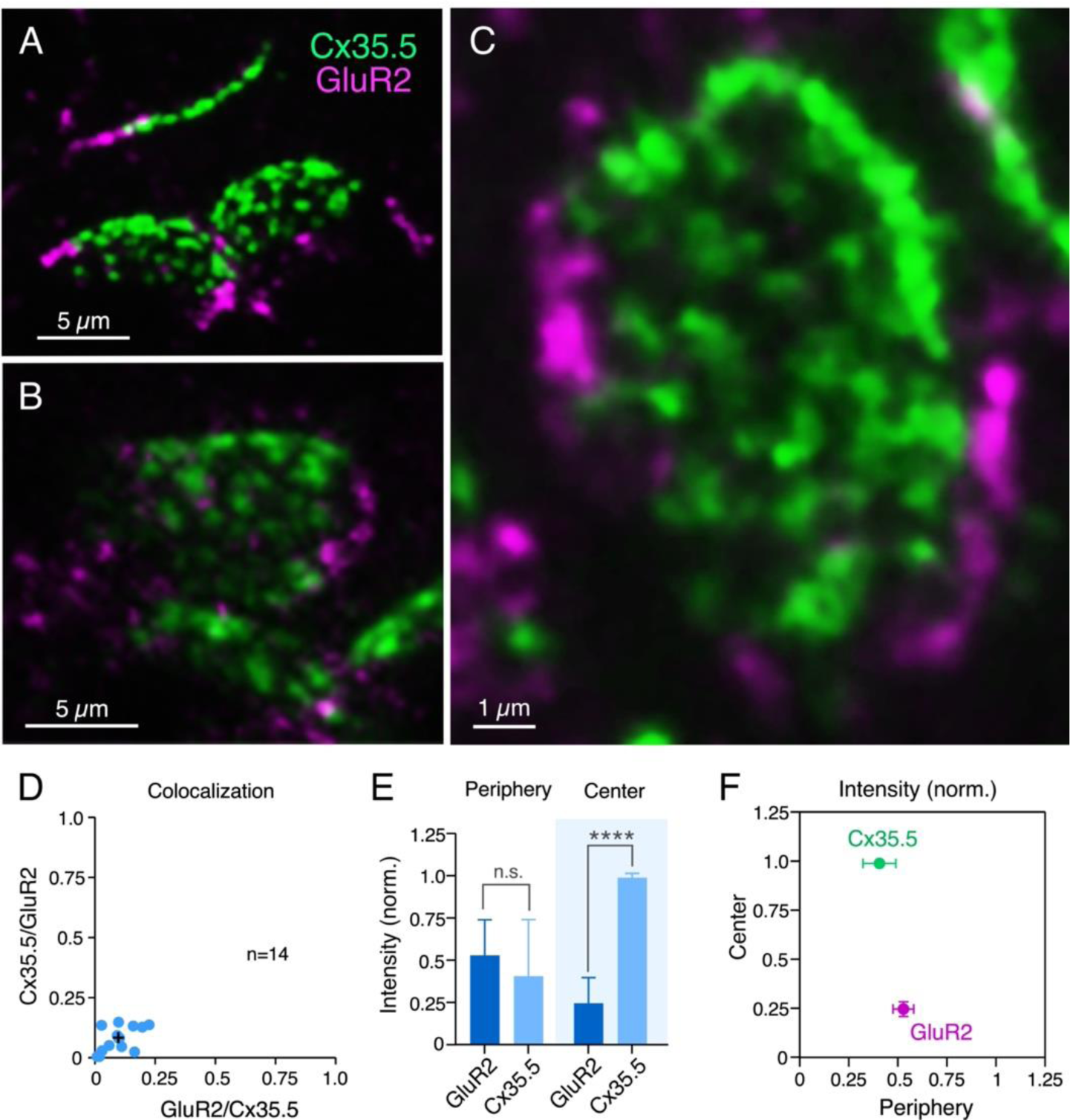
Electrical and chemical transmitting areas are mutually exclusive. (A,B) Expanded synaptic contact areas labeled with anti-Cx35.5 (green) and anti-GluR2 (magenta). (C) ‘En face’ view of an expanded synaptic contact area showing that GluR2 labeling is restricted to the periphery of the contact, whereas Cx35.5 labeling is distributed throughout the whole contact area. (D) Graph shows the lack of colocalization between Cx35.5 and GluR2 fluorescence at individual CE contacts, determined by the Mander’s Colocalization Coefficient: GluR2/Cx35.5 0.078 ± 0.05 (x axis); Cx35.5/GluR2 0.098 ± 0.06 (y axis), n= 14 (mean ± SEM). (E) Distribution of labeling. Quantification of labeling for Cx35.5 and GluR2 at the periphery and the center of the CE contact area (Student’s t-test <0.05, n=4 on face views, p=<0.0001). Error bars denote ± SEM. GluR2 fluorescence is higher in the periphery, while Cx35.5 fluorescence is significantly higher at the center of the contact area. (F) Graphical description of the center vs. periphery distribution of Cx35.5 and GluR2 for the data described in panel E.

Rather than diffusely distributed, labeling for Cx35.5 in expansion samples was characterized by well-defined puncta distributed throughout the contact area, suggesting that they each might represent an individual GJ plaque. Consistent with this interpretation, double labeling for Cx35.5 and Cx34.1 (Fig, 3A,C) showed a high degree of colocalization at CE contact areas (Fig 3E). A high degree of colocalization (Fig. 3B,D) was also found between labeling for Cx35.5 and ZO1 (Fig 3F). Moreover, labeling for Cx35.5 and Cx34.1 colocalized at single puncta, as indicated by line scan of individual puncta. Given the characteristic concavity of the CE contact area, the relationship between labeling for pre- and postsynaptic GJ proteins (Fig. 4A) was easier to establish and accurately measure at the periphery of the labeled areas (Fig. 3B). Line scan of samples labeled for Cx35.5 and Cx34.1 indicated the presence of labeling for these proteins at single punctum (Fig. 4C). Strikingly, while the peak of maximum intensity for the pre-and postsynaptic connexins were aligned (Fig. 4C), the peak of maximum intensity for Cx35.5 and ZO1 were distanced (Fig. 4D). This is consistent with the fact that while Cx35.5 is presynaptically localized, ZO1b, one of the two zebrafish orthologs of ZO1, was reported to be postsynaptic. The observed distance between the peaks of fluorescence was not due to the differences in the wavelength of the fluorophores, and it remained when secondary antibodies were inverted (Fig. 4D-E). These differences are quantified in the graph of Figure 4F. The finding confirms previous conclusions reached with biochemical and chimera analysis indicating that ZO1b is only located at postsynaptic hemiplaques (12). Although a distance between Cx35.5 and Cx34.1 labeling peaks was observed in some examples, it was still too small to be detected with this method due to the spatial amplification produced by the fluorophores. Thus, by increasing the distance between pre- and postsynaptic sites, expansion increases the resolution, making it possible to explore differences in the composition of GJ hemiplaques. Altogether, our findings indicate that each puncta correlates to a GJ.

**Figure 3.**
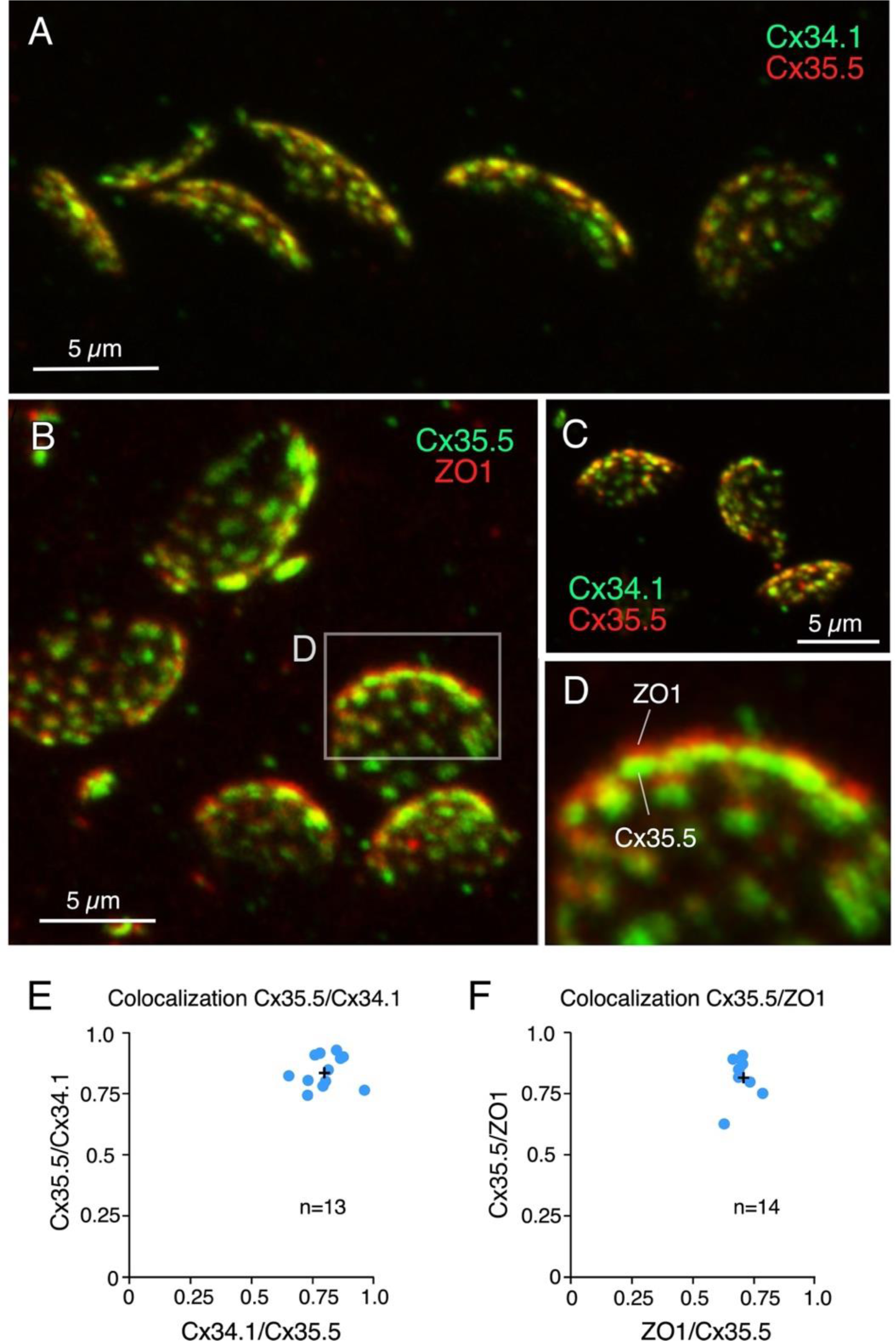
Labeling for gap junction proteins reveals the presence of multiple puncta at expanded CE synaptic contact areas. (A) CE synaptic contact areas labeled with anti-Cx34.1 and anti-Cx35.5. (B) Contact areas labeled with anti-Cx35.5 and anti-ZO1. Same experiment as Fig. 1E. (C) Labeling with anti-Cx34.1 and anti-Cx35.5. (D) Magnification of the boxed region in panel B showing a side view of an expanded synaptic contact area labeled for Cx35.5 and ZO1. (E) Graph showing colocalization of Cx35.5 and Cx34.1 fluorescence at individual CEs determined by the Mander’s Coefficient: Cx34.1/Cx35.5 0.80 ± 0.08 (x axis); Cx35.5/Cx34.1 0.84 ± 0.06 (y axis), n=14. (F) Colocalization of Cx35.5 and ZO1 fluorescence at individual CEs. Mander’s Coefficient: ZO1/Cx35.5 0.71 ± 0.04 (x axis); Cx35.5/ZO1 0.82 ± 0.07 (y axis), n=14.

**Figure 4.**
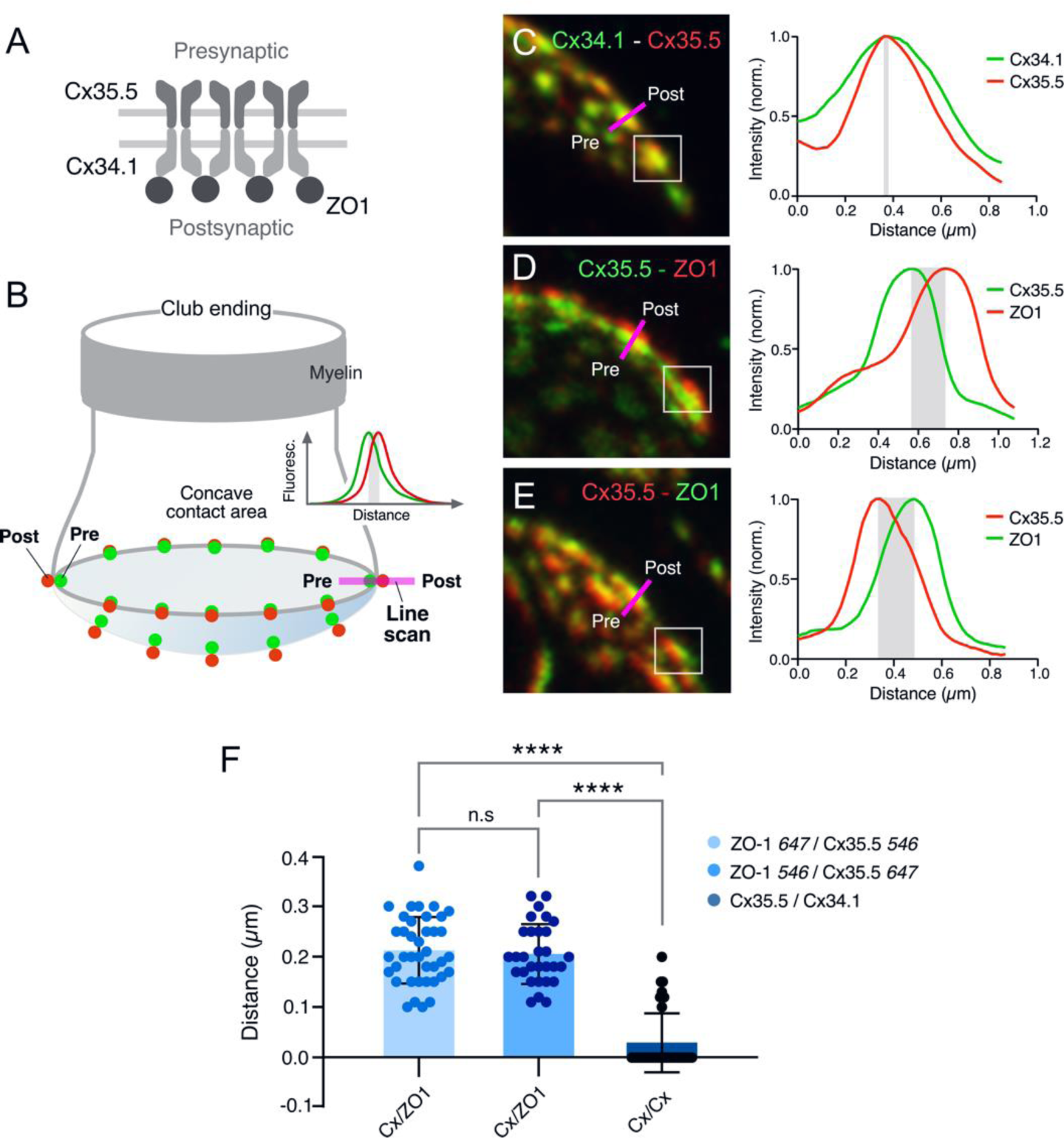
Expansion microscopy reveals the molecular components of gap junction plaques at CE synaptic contact areas. (A) Schematic representation of the molecular organization of GJs between CEs (pre-synaptic) and the M-cell (post-synaptic). The presynaptic and postsynaptic hemichannels are formed by Cx35.5 and Cx34.1, respectively. The scaffolding protein ZO1 is postsynaptic and interacts with Cx34.1. (B) Cartoon of a CE terminal illustrating the concavity of its contact area with the M-cell. The concavity determines differences in the relative position of presynaptic (green) vs. postsynaptic (red) labeling at different points throughout the contact area. Puncta located in the periphery of the contact are ideally aligned to determine colocalization of fluorescence at individual puncta (line scan, inset). (C-E) Line scan of puncta at expanded contact areas showing colocalization of presynaptic Cx35.5 and postsynaptic Cx34.1 (C), and presynaptic Cx35.5 and ZO1 (D-E). The example in C is part of the experiment illustrated in Fig. 3A. The magenta lines indicate the position of the line scan in each case. The fluorescence intensity profiles for each fluorophore are illustrated on the right side of each panel. As a control, secondary antibodies were swapped in E. (F) Bar graph illustrates the distance between the peaks of fluorescence intensity profiles for Cx35.5-Cx34.1 labeling (either 647Atto or 546Alexa-Cx35.5 vs. either 647Atto or 546Alexa -Cx34.1: 0.03 ± 0.001 µm, n=38) and Cx35.5-ZO1 labeling (546Alexa-ZO1 vs. 647Atto-35.5: 0.21 ± 0.002 µm, n=39). Secondary antibodies were swapped as control (647Atto-ZO1 vs. 546Alexa-Cx35.5: 0.20 ± 0.001 µm, n=30). Bars represent ± SEM. Anova analysis with Tukey’s multiple comparison test correction p<0.001.

Puncta labeled for Cx35.5 (Fig. 5A-B), which we established each represent a GJ, showed a high degree of variability in number and size (see inset of Fig. 5C). We then quantified the number of puncta and their individual area in ‘En face’ views of CEs, at which this analysis was possible. [Panels B and C in Fig. 5 are the same experiment as Figs. 1E and 3B-C, illustrating the ability of expansion microscopy for providing multiple layers of information within the same experiment.] Variability in the number and area of puncta was observed between neighboring CEs within the same M-cell lateral dendrite (Fig. 5D). The number of GJs per CE averaged 36.72 ± 0.4 (n=11), and their area ranged from 0.05 to 1.98 µm^2^. The number of GJs might have been slightly underestimated because fluorophore spatial amplification could have caused a big punctum to form from two closely spaced small GJs. This limitation could result in two nearby GJs appearing to merge. After correcting for the expansion factor (13×) and assuming that connexons in GJ plaques are organized in a crystalline fashion with a density of 12,000 connexons/µm^2^ (38), we estimated that GJs at CEs contain 49 to 1775 connexons, and the total junctional area represents an average of 12,706.6 ± 129.1 (n=11) connexons per CE. Thus, electrical transmission at zebrafish CEs is mediated by multiple GJs containing a variable number of channels.

**Figure 5.**
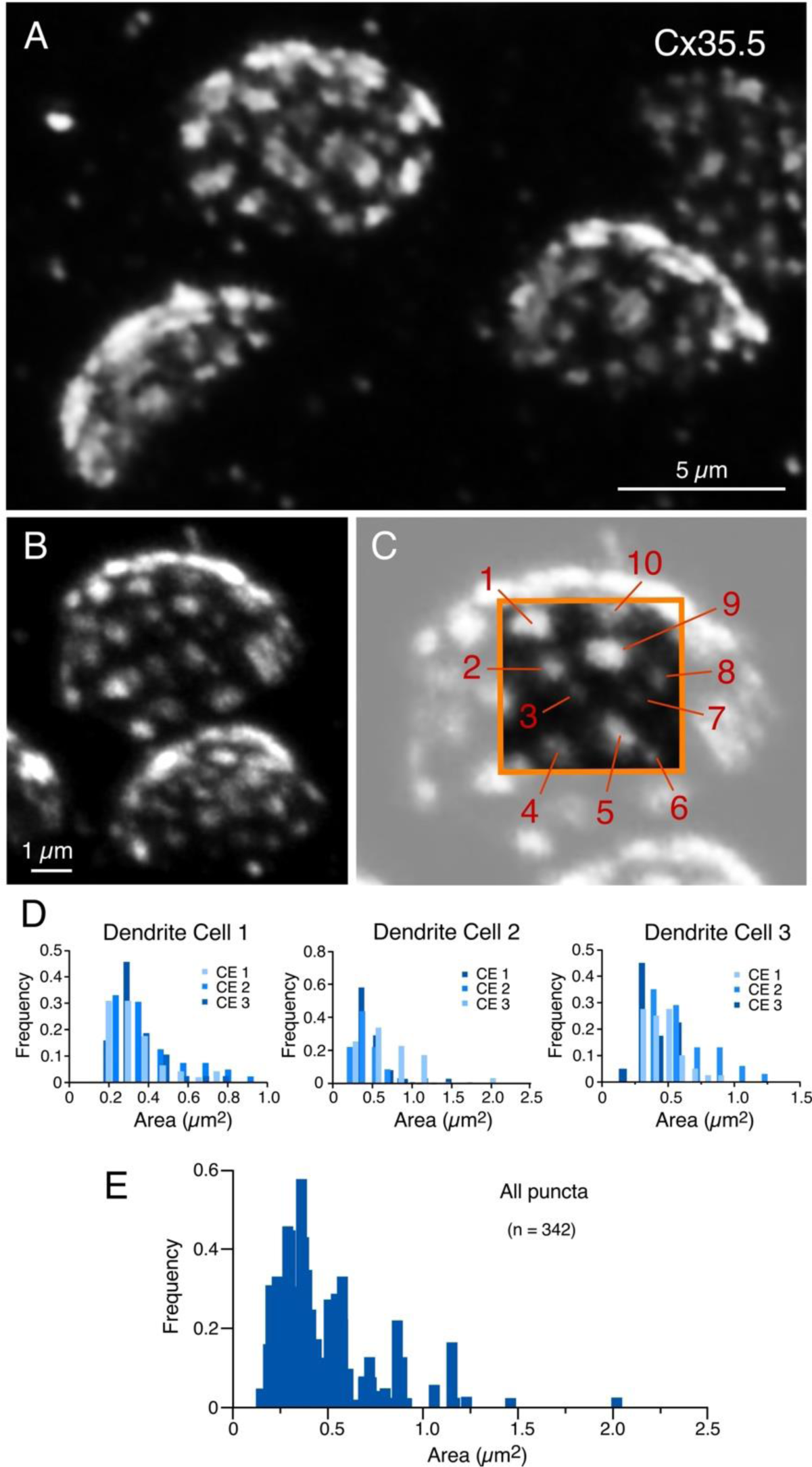
Expansion reveals the presence of multiple, variably sized, gap junctions. (A-B) ‘En face’ views of expanded CE contact areas labeled with anti-Cx35.5 showing multiple puncta with high variability of their size. (C) Magnification of the CE contact area at the top of panel B (light gray). The area enclosed by the orange box illustrates the wide variability in puncta size, labeled 1 to 10 (with the goal of highlighting puncta size variation, the image delimited by the orange box was cropped from and placed on the same region of the lighter image). Panels B and C are the same experiment as Figs. 1E and 3B-C, demonstrating the ability of expansion microscopy for providing multiple layers of information in the same experiment. (D) Frequency histograms illustrating the number and size distribution of puncta labeled for Cx35.5 within individual CEs. Histogram shows the distribution of puncta size at three CEs obtained in the same cell (dendrite), illustrating overlap in different shades of blue. Three different examples are illustrated from left to right. Each histogram shows similar variability in number and size for all nine terminals. (E) Frequency histogram of number and size distribution of Cx35.5 puncta for the nine reconstructed contacts in panel D.

Our labeling with Cx35.5 and GluR2 showed that most of the contact area of a CE operates as an electrical synapse. We then asked what other anatomical structures might be contributing to electrical transmission. Previous electron microscopy analysis exposed the presence of adherens junctions (AJs) in close proximity to neuronal GJs, including those at CEs in goldfish and larval zebrafish (Fig. 6A). AJs are known to initiate and mediate the maturation and maintenance of cell-cell contacts, including GJs, at which a wealth of evidence suggests a close functional interaction (39–41). Therefore, we decided to investigate the incidence, association, and spatial distribution between these two structures at CEs. Recent immunohistochemical analysis revealed the association between components of AJs and Cx36 in various mammalian structures (42). Expansion for Cx35.5, transmembrane protein N-cadherin, and the intracellular protein ß-catenin, the latter two being major structural components of AJs, showed that labeling for AJs components seem equally distributed through the synaptic contact (Fig. 6B-D). In contrast to Cx35.5, which is characterized by well-defined puncta reminiscent of GJs, labeling for N-cadherin and ß-catenin was diffuse, less structured, and did not colocalize with Cx35.5 (Fig. 6B). Rather, the labeling for N-cadherin (Fig. 6C) and ß-catenin (Fig. 6E) was interleaved with Cx35.5 puncta, as illustrated by line scan analysis in the margins of CE contacts (Figs. 6D and 6F). Furthermore, in ‘En face’ views, labeling for N-cadherin (Fig. 6G) and ß-catenin (Fig. 6H) was broadly distributed throughout the surface of the contact, appearing to engulph Cx35.5 labeled puncta. While labeling for N-cadherin and ß-catenin was interleaved with Cx35 labeling, we observed some degree of colocalization (Fig. 6B) likely due to fluorophore amplification of the close spatial association between GJs and AJs (see Fig. 6A).

**Figure 6.**
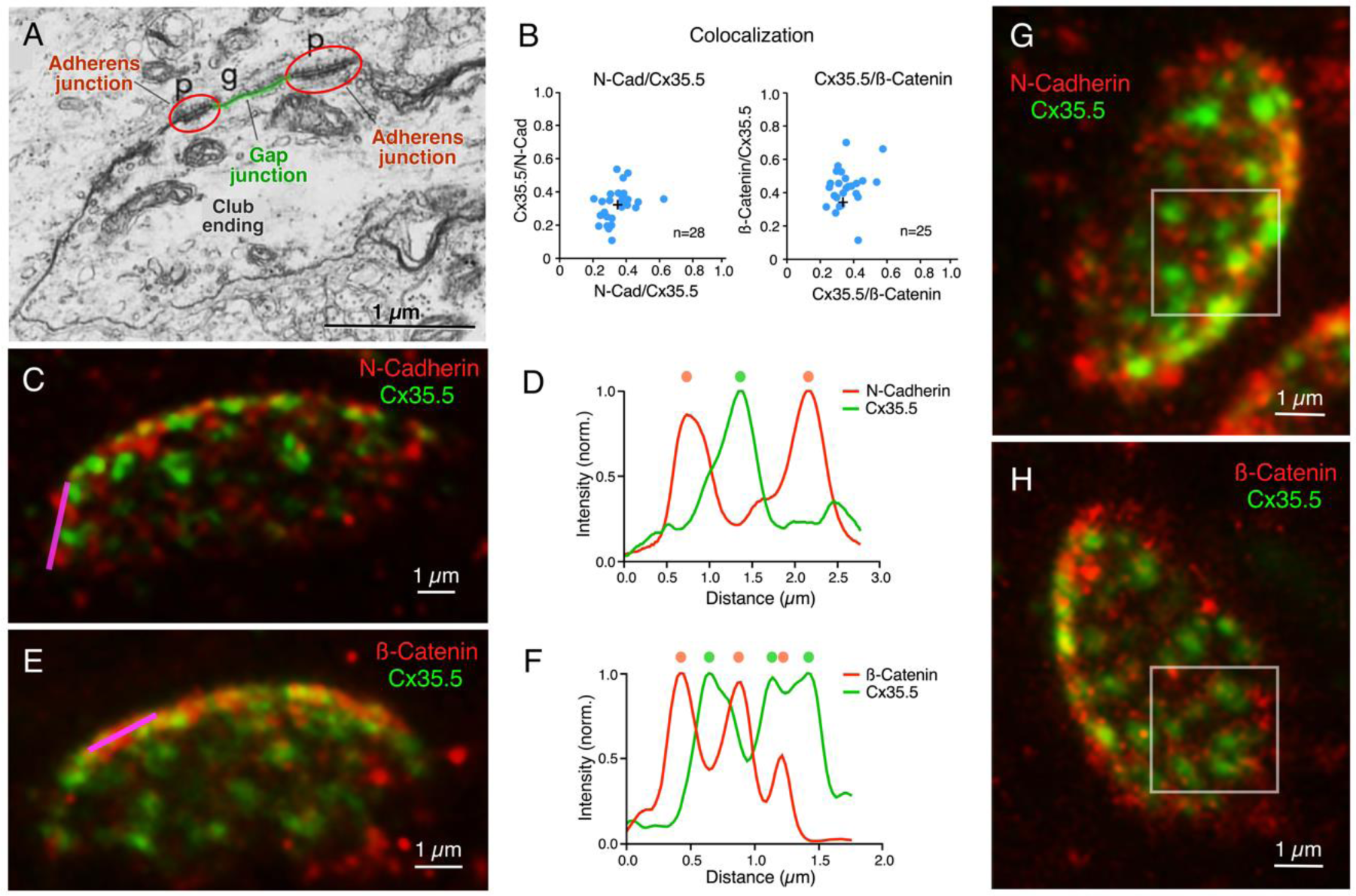
Gap junctions at CEs are associated with adherens junctions. (A) Electron micrograph of a CE obtained in a 6 dpf zebrafish showing a GJ (g, highlighted in green) surrounded by AJs (p, encircled in red). From Kimmel et al., 1981 (with permission) (49). (B) GJ and AJ proteins do not colocalize. Left: Cx35.5 and N-Cadherin labeling show a low index of colocalization. Mander’s coefficient: N-Cadherin/Cx35.5 0.35 ± 0.003 (x axis); Cx35.5/N-Cadherin 0.32 ± 0.003 (y axis), n=28. Right: Cx35.5 and β-catenin labeling also show low colocalization. Mander’s coefficient: Cx35.5/β-catenin 0.43 ± 0.004 (x axis); β-catenin/Cx35.5 0.36 ± 0.008 (y axis), n=25. (C) Expansion microscopy of a CE contact area labeled for N-cadherin (red) and Cx35.5 (green). (D) Fluorescence intensity profiles for N-cadherin and Cx35.5 obtained with a line scan (magenta line in C) are mutually exclusive. The low degree of colocalization observed in panel B is likely due to fluorophore amplification and the close spatial association between GJs and AJs, as shown in panel A. (E) Image shows an expanded CE contact area labeled for β-Catenin (red) and Cx35.5 (green). (F) Line scan (magenta line in E) shows that labeling for β-Catenin and Cx35.5 are also mutually exclusive. (G,H) ‘En face’ view of the expanded contact area double-labeled for N-Cadherin and Cx35.5, and β-Catenin and Cx35.5, respectively, showing the close association of GJs and AJs throughout the synaptic contact area. Insets: the boxed areas in the ‘En face’ images highlight the mutually exclusive labeling.

The spatial distribution of the labeling for components of AJs suggest that their function is associated with electrical transmission, rather than chemical transmission. To establish this association, we measured the relative proportion of labeling fluorescence of various synaptic components at CE contact areas. As expected, labeling for Cx35.5 and the GJ scaffold ZO1 were equally proportional (Fig. 7A). Similar proportionality was found for fluorescence of N-cadherin and Cx35.5, and ß-catenin and Cx35.5 (Fig. 7B). In contrast, fluorescence of GluR2 represented a smaller fraction than that of Cx35.5 labeling (Fig. 7C). Finally, we normalized the fluorescence to the area of the CE contact to calculate the area occupancy for these three synaptic structures (Fig. 7D). Less than 20% was occupied by GluR2, while 60% of the contact was occupied by equal amounts of Cx35.5 and AJs. About 20% of the contact area was not labeled for these antibodies, either corresponding to the surface membrane between the structures recognized by labeling, or to the presence of additional synaptic components. Thus, while chemical synapses are restricted to a small and peripheral area of the contact, most of the contact surface is occupied by multiple GJs of variable size which are interleaved and closely associated to AJs (Fig. 7E).

**Figure 7.**
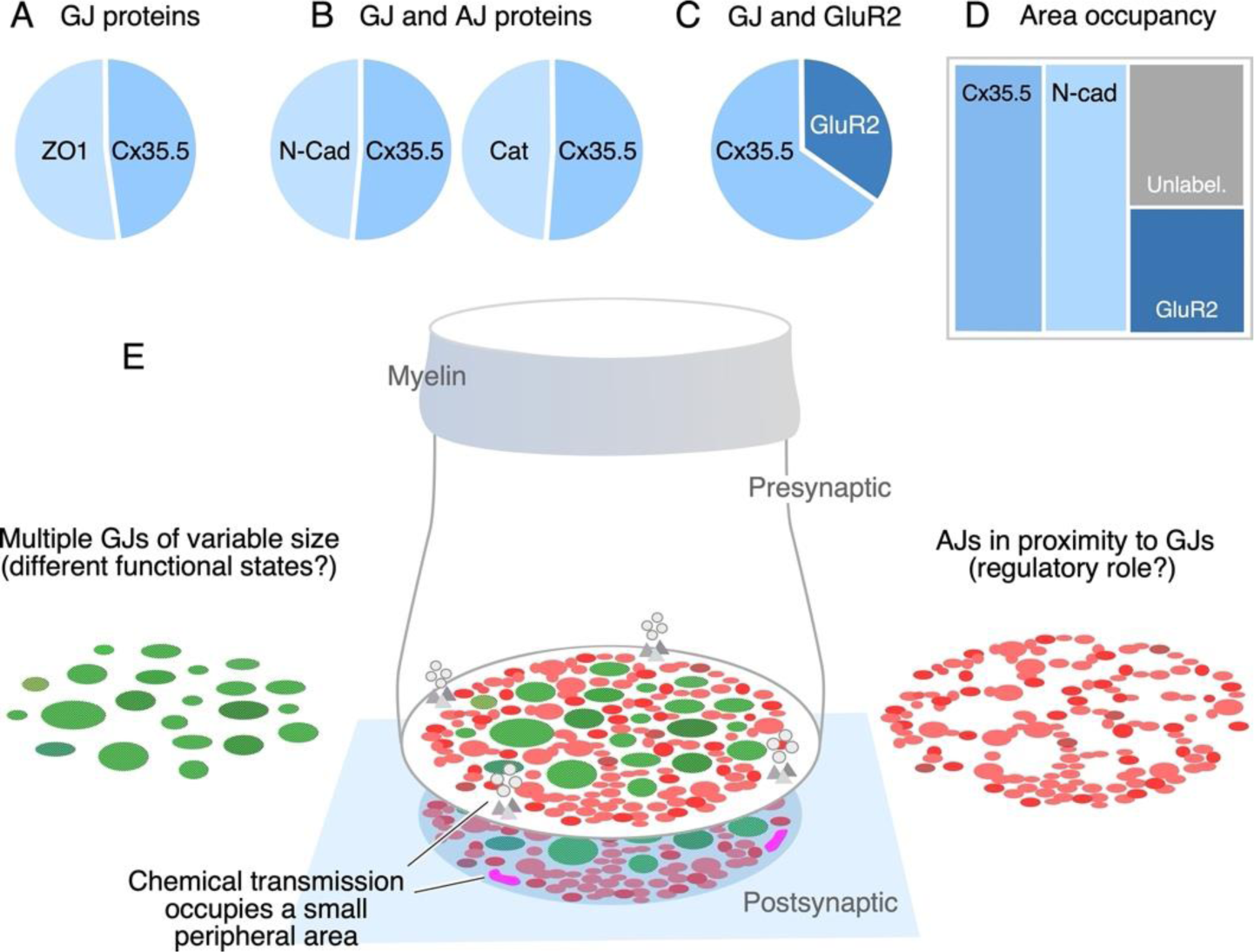
The electrical synapse at the CE combines multiple gap junctions with adherens junctions. (A) Double labeling with anti-Cx35.5 and anti-ZO1 show similar proportion of fluorescence at CEs (ZO1 43.3% ± 1.49; Cx35.5 38.25% ± 1.49, n=8). (B) Double labeling for N-Cadherin and Cx35.5 (left), and for β-Catenin and Cx35 (right) also show similar proportionality (N-Cad 41.88% ± 0.82, n=13; β-Catenin 39.84% ± 0.49; Cx35.5 38.25% ± 0.59, n=11). (C) Double labeling for Cx35.5 and GluR2 shows lack of proportionality, with Cx35.5 fluorescence occupying the majority of the CE contact area (GluR2 18.9% ± 0.79; Cx35.5 35.91% ± 1.25; n=11). (D) Tree plot illustrating the area occupancy (fluorescence per contact area) for AJ (N-Cad=29.1%), GJ (Cx35=30.8%), and glutamatergic (GluR2=18.9%) components at individual CEs. The unlabeled area represents 21.3% of the contact’s surface. (E) The cartoon summarizes the synaptic components identified at a single CE contact. While chemical synapses are restricted to a small and peripheral area of the contact (presynaptic vesicles and release sites are represented in grey, postsynaptic receptor areas in magenta), most of its contact surface is occupied by multiple gap junctions (GJs; green) of variable size, which are interleaved and closely associated to adherens junctions (AJs; red).

## DISCUSSION

> *“One of the most difficult problems in the correlation of structural and functional concepts is that of accurately defining the synapse.” J. David Robertson (1963)* (26)

Because they support both electrical and chemical synaptic transmission, CEs on the M-cells enabled the correlation of their synaptic properties with structural specializations for each of these modalities of communication (26). In addition to specializations observed at purely chemically-transmitting contacts, electron microscopy analysis by Robertson revealed areas of close membrane apposition, which in ‘En face’ views were oval-shaped and exhibited a characteristic reticular pattern (43). These ‘synaptic discs’ provided early evidence for the cellular structures that we now know as ‘gap junctions’ (GJs): clusters of intercellular channels which provide the mechanism of communication for electrical transmission. However, in his seminal paper (26), Robertson warned about the limitations of correlating structural and functional notions to define the components of an electrical synapse. In contrast to chemical synapses, it is challenging to define exactly what anatomically constitutes an electrical synapse, as neuronal GJs are often found connecting cell somata or other neuronal processes, such as dendrites and axons (44). Because the presynaptic bouton anatomically marks the limit of a synapse, CEs provided the ideal opportunity to investigate the components of an electrical synapse. By applying expansion microscopy to these single terminals, we found evidence suggesting that, in contrast to the general perception, electrical synapses might have a more complex structural organization. Our data indicates that the CE electrical synapse operates with, not one, but multiple GJs that are in close association with structural and signaling molecules known to be components of AJs (Fig. 7E). This extended overarching notion of the synaptic organization of an electrical synapse might contribute to the understanding of their diversity of functional and structural organization.

### Expansion microscopy of a single synaptic contact

Because of their identifiability, CEs have historically been amenable for exploring synaptic structure and function with novel technical approaches. We applied expansion microscopy to these terminals, which allowed us to generate a map of the distribution of its various synaptic proteins. While the sentence, “The map is not the territory,” was coined by Alfred Korzybski to metaphorically illustrate the distinction between brain perception and reality (45), it applies to the properties of maps in general and to the usefulness of applying expansion microscopy to expose the synaptic map of CE contacts. The properties of a map are determined by the approach used to extract the information that generates it. Moreover, any useful map should not necessarily be fully accurate (discussed in “Of Exactitude in Science” by Jorge Luis Borges (46)), but capable of capturing an element of reality. As a representation of reality, the labeling of molecules forming GJs, glutamate receptors and AJs, allowed us to expose the incidence and spatial distribution of the various synaptic components with enough accuracy to provide an ‘analogous structure’ of the contact’s structural organization. We found that most of the contact area of a CE operates as an electrical synapse and only a peripheral ∼20% is dedicated to chemical transmission. The 4× diameter increase after expansion represented a 13× increase of the oval contacts areas and did not alter the morphology of the terminals, as supported by: 1) expansion reproduced the characteristic concavity of the CE synaptic contact area with the M-cell; 2) the spatial distribution of chemical vs electrical synaptic areas was consistent with that observed by electron microscopy in adult goldfish (36) and unexpanded zebrafish terminals (12); 3) each of the labeled puncta contain all the molecular components known to form GJs at CEs; 4) morphometric analysis indicated that the expansion process had no selective effects across the CE population; 5) Finally, tissue expansion was tridimensional and exposed the expected pre-vs postsynaptic localization of proteins at single punctum representing individual GJs. Expansion microscopy is complementary to a different representation of reality, electron microscopy, which is capable of providing fine details of the structures, but can inform, to a much less degree, their biochemical composition. Unlike the more labor-intensive electron microscopy, expansion microscopy was more easily capable of reconstructing the complete contact area of the terminal, while additionally exposing its biochemical composition. Thus, without having its structural resolution, expansion microscopy was able to expose, with sufficient resolution, spatial features that were only observed so far with electron microscopy.

Furthermore, because of its tridimensional nature, expansion microscopy not only enlarged the synaptic contact area, but also the distances between synaptic elements, enhancing the detection of pre-vs postsynaptic components at GJs. Ultrastructural images and the bilateral requirement of GJ hemichannels convey the sense that GJs are symmetrical structures. Emerging evidence obtained at CEs suggest otherwise. Recent data indicates that GJ asymmetry at CEs is not only restricted to the connexin composition of pre- and postsynaptic hemichannels, but also to scaffolding molecules (12, 14, 31, 47, 48). That is, biochemical data and chimera analysis suggested that ZO1b, one of the two zebrafish orthologs of ZO1 whose function is critical for the presence of connexins at electrical synapses, is located in the postsynaptic hemiplaque from where it exerts its critical function (12). Confirming this conclusion, our expansion microscopy unambiguously exposed the postsynaptic location of ZO1b. Moreover, because the antibody recognizes both ZO1b and ZO1a, our data indicates that, although with different functions, both ZO1 orthologs are restricted to postsynaptic hemiplaques.

### Functional association between gap junction and adherens junctions

Electron microscopy of CEs in both adult goldfish and larval zebrafish revealed the presence of AJs in the vicinity of GJs (35, 49), which was also observed at neuronal GJs in viper cerebellum (50) and cat inferior olive (51). AJs are dynamic cell-cell adhesion complexes that are continuously assembled and disassembled. They are formed by a complex of proteins that include cadherins and catenins (ß and alfa). Growing evidence indicates a functional relationship between AJs and GJs (39, 52–55). While it was generally accepted that GJs form via lateral diffusion of hemichannels following microtubule-mediated delivery to the plasma membrane, more recent evidence obtained in cell expression systems shows that microtubules actually tether to the AJ, facilitating delivery of vesicles containing connexin hemichannels directly to the cell-cell border of the GJ (39). This process depends on the interaction of microtubules with plus-end-tracking protein (+TIP) EB1, its interacting protein p150 (Glued), and the AJ proteins N-cadherin and ß-catenin (39). Furthermore, the evidence indicates that this process also requires homophilic interactions between N-cadherins. Similar peripheral delivery of GJ channels to AJs was observed in sensory epithelial cells of the cochlea (40), indicating that this mechanism also operates in tissues. Also consistent with a close functional relationship between these structures, AJ formation hierarchically regulates the formation of GJs in cardiac pacemaker cells (41), adjusting the excitability and coupling of these neurons in the context of their pacemaking function.

### The components of an electrical synapse

The investigation of the mechanisms and structures underlying electrical transmission has generally centered on the properties of GJ channels and their supporting molecules, a perspective restricted to a single GJ plaque. However, electrical transmission at each zebrafish CE terminal is mediated by about 30 GJs, suggesting that multiple GJs operate as a functional unit. Like chemical synapses that operate with dramatically different numbers of releases sites, ranging from a single one at synaptic terminals of PHP cells on the M-cell (56, 57) to up to 700 in the Calyx of Held (58, 59), electrical synapses could function with different number of GJs. Because of the wide variation in size, GJs at CEs might coexist at different states of conductance. Operating with multiple GJs might result in more reliable electrical transmission, as only a very small fraction of channels are known to be functional at individual GJs (28, 60–62). The presence of multiple GJs might also be indicative of high plastic regulation, a possibility consistent with the dynamic properties of CE electrical synapses, which are known to undergo activity-dependent potentiation of their synaptic strength (63–66). While it is presently unknown how regulation of the overall conductance of the CE electrical synapse is achieved, it must necessarily result from the coordinated contribution of its multiple GJs, suggesting the need of additional synaptic structures capable of coordinating the function of the numerous GJs. Given the close spatial association that we report here, it is tempting to speculate that interactions between AJs and GJs could underlie such function. A close association between components of AJs and Cx36 was observed at various mammalian brain structures (42), and it was suggested they share a common molecular complex (42). Since interactions between GJs and AJs involving cadherins are thought to be relevant for the insertion of new GJ channels (40, 67), this process might be intimately related to the maintenance and plasticity of GJ communication via regulated turnover of these channels at CEs, which are formed by fish homologs of Cx36 (33, 68). Consistent with the possibility of AJs promoting the insertion of new GJ channels at CEs, microfilaments arriving to an AJ situated in close proximity to a GJ can be observed in an image of the EM study by Kohno and Naguchi; see Fig. 2 in Ref. (35). Operating with multiple GJs also provide electrical synapses with the possibility of either silencing or activating individual GJs as a potential additional mechanism of strength regulation. Finally, although N-cadherin was also shown to be involved in regulating chemical synapses (57, 83), the distribution of chemical synapses to small peripheral areas of the CE contact indicates a primary functional role of this molecule at electrical synapses, as both GJ proteins and N-cadherin are similarly distributed throughout the entire synaptic contact area. Future investigations on the functional association between GJs and AJs will shed light on the functional organization of electrical synapses.

The structural complexity of chemical synapses has long been recognized. Structural complexity is also a hallmark of immunological synapses (69), specialized functional contacts which quickly assemble between a thymus-derived lymphocyte and an antigen presenting cell (70, 71). Both chemical and immunological synapses share a general molecular organization combining cell-adhesion molecules, usually restricted to the periphery of the contacts, with those more centrally located and responsible for providing intercellular communication (72). Our results indicate that this structural arrangement might also apply to the CE electrical synapse. That is, our finding that AJs surround GJs all through the contact is consistent with the general organization of chemical and immunological synapses, at which adhesion molecules are located in the periphery of the communicating mechanism. Thus, just as chemical and immunological synapses, electrical synapses also seem to combine cell adhesion molecules with those responsible for mediating intercellular communication. Based on these similarities in the overall organization across synapses and evidence of functional interactions with GJs (see above), we propose that AJs could be considered components of electrical synapses. While additional components are likely to contribute to electrical transmission, the distribution of adhesion molecules at electrical synapses might serve to define their synaptic boundary, an arrangement that might be used as a template to identify electrical synapse elsewhere. Accordingly, multiple Cx36-labeled puncta engulphed by N-cadherin labeling was also observed at electrical synapses between the somata of neurons of the mesencephalic nucleus of the trigeminus (see Fig. 8 in Ref. (42)), indicating that this arrangement is not a unique feature of mixed synapses but might also apply to electrical synapses elsewhere.

Thus, as single synapses, expansion microscopy of CEs offered the possibility of identifying the structures that support electrical transmission and, therefore, the components that define an electrical synapse, which can be used to define electrical synapses throughout animal connectomes. Moreover, defining the components of an electrical synapse will help to expose their functional diversity, as different synapses might operate with different numbers of GJs and synaptic arrangements. Together with the genetic accessibility of zebrafish, the synaptic map provided by expansion microscopy will facilitate exploring the functional relationship between the structures supporting electrical transmission and its regulation.

## METHODS

### Experimental model and subject details

All experiments were performed in 5 days post fertilization (dpf) zebrafish, *Danio rerio*. Zebrafish were housed at the Department of Neuroscience zebrafish facility, bred and maintained at 28°C on a 14-hour light / 10-hour dark cycle. Experiments were carried out in the Tol-056 enhancer trap line (73).

### Immunohistochemistry

Zebrafish larvae were anesthetized with 0.03% MS-222 (tricaine methanesulphonate) and fixed for 3 hours with 2% trichloroacetic acid in 1X PBS. Fixed samples were then washed 3 times with 1X PBS, followed by a brain dissection with the help of a custom-made tungsten needle and forceps. The needle was also used to remove the cerebellum, optic tectum, and telencephalon to accommodate for the working distance of the microscope objective. This step was not necessary in samples used for expansion methods. The dissected brains were washed with 1X PBS + 0.5% Triton X-100 (PBS-Trx) and blocked with 10% Normal Goat Serum + 1% DMSO in PBS-Trx. Brains were then incubated with a primary antibody mix at room temperature overnight. The antibody mix, with block solution, included combinations of the following: rabbit anti-Cx35.5 (Miller et al., 2015 (74), clone 12H5, 1:200), mouse IgG1 anti-Cx35/36 (Millipore, cat. MAB3045, 1:250), mouse IgG2A anti-Cx34.1 (Miller et al., 2015 (74), clone 5C10A, 1:200), mouse IgG1 anti-ZO1 (Invitrogen, cat. 33–9100, 1:200), mouse IgG1 anti-N-cadherin (BD Transduction Laboratories, cat. 610920, 1:50), mouse IgG1 anti-Beta-catenin (Sigma, cat. C7207, 1:100), rabbit IgG anti-GluR2/3 (Millipore, cat. 07-598, 1:200), and chicken IgY anti-GFP (Abcam, cat. ab13970, 1:200). After 3 washes in PBS-Trx, brains were blocked with 10% Normal Goat Serum + 1% DMSO in PBS-Trx, followed by incubation in secondary antibody mix at room temperature for 4 hours. Secondary antibody mixes included, in addition to block solution, combinations of mouse IgG-Alexa Fluor 546 (Invitrogen, cat. A11030, 1:200), mouse IgG-Alexa Fluor 647 (Invitrogen, cat. A21235, 1:200), rabbit IgG-Alexa Fluor 546 (Invitrogen, cat. A11010, 1:200), rabbit IgG-Atto 647N (Sigma-Aldrich, cat. 40839, 1:200), mouse IgG-Atto 647N (Sigma-Aldrich, cat. 50185, 1:200), and chicken IgY-Alexa Fluor 488 (Invitrogen, cat. A11039, 1:200). Samples were washed with 1X PBS 4 times and overnight at 4°C, and then transferred in the dark, onto a slide, and mounted with ProLong Gold antifade (Invitrogen, cat. P36930). Finally, samples were covered using the ‘bridge’ procedure (75) and sealed with nail polish.

### Expansion

Expansion of brain samples was performed following previous protocols (Asano S, *et al*, 2020) with additional modifications, using the following reagents: Anchoring solution: Acryloyl-X SE (10mg/ml); Monomer solution: 50% acrylamide, 2g/100ml N,N’ methylenebisacrylamide, 5M NaCl, 10X PBS, 38% Sodium Acrylate, and ddH2O to a complete volume; Inhibitor solution: 4-hydroxy-TEMPO (4-HT) (0.5%); Accelerator: Tetramethylethylenediamine (TEMED) (10%); Initiator: Ammonium Persulfate (APS) (10%); Digestion buffer: 1M Tris (pH 8.0), 0.5 M EDTA, 100-X Triton, 5M NaCl, and ddH2O to a complete volume, and Proteinase K (8 units/mL). Upon completion of immunolabeling steps, the sample was incubated in anchoring solution (1:100) overnight. The sample was then washed twice in 1X PBS and placed into the following mix: monomer solution, inhibitor solution, the accelerator (TEMED), and the initiator (APS), and placed in 4°C for one hour, and then at 37°C for two hours. Once the hydrogel polymerized the sample was placed for 16 hours at 50°C in the digestion buffer + Proteinase K. Following this procedure, the hydrogels were imaged with the confocal microscope.

### Confocal imaging

All images were acquired on LSM 710 and LSM 880 Zeiss microscopes using the following laser wavelengths: Argon 458/488/514, HeNe 543, and HeNe 633, along with the corresponding filter: MBS 488/543/633, MBS 458/543, MBS 488/543/633, using either a 40× 1.0 NA water immersion objective or a 63× 1.40 NA oil immersion objective. Confocal settings were adjusted to achieve maximum visualization of labeling at CE contact areas. Both lateral and ‘En face’ views of expanded CE contact areas were used for quantitative analysis.

### Quantification and statistical analysis

Confocal images were obtained using ZEN (black edition) software and analyzed with FIJI. Contrast and brightness of fluorescence channels were individually adjusted and contrasted in Photoshop (Adobe), using blur and sharpen filters. Nonetheless, quantitative analysis of image fluorescence was carried out using raw data. For quantification of Cx35 puncta area, each punctum was defined as a region of interest (ROI) and then its area calculated. To compare the center/periphery distribution of Cx35.5 and GluR2 staining, each CE contact area was defined as an ROI and divided into a peripheral ring (∼1/4 of area) and a center area (∼3/4 of area). Fluorescence intensity for each labeling at each area of the ROI was then compared. For statistical comparison, fluorescence was normalized to account for the variability between the samples. Fluorescence colocalization was estimated with the Mander’s coefficient (“JaCoP” plugin, FIJI) once the CE contact area was defined as a ROI. For estimates of area occupancy of each labeling, the area of the synaptic contact was estimated by combining Cx35.5 and GLUR2 labeling, which together constitute a good approximation of the contact’s surface area. For statistical analysis, unpaired Student t-test or one-way ANOVA with Tukey’s multiple comparsions test to compare multiple groups were performed using GraphPad Prism software. Data is presented as the mean ± standard error of the mean (SEM).

## Acknowledgments

We thank Martin Pinter, Anne Martin, Adam Miller, and members of the Pereda lab for critical feedback on the work and manuscript. We also thank Martin Pinter for his help on the statistical analysis of the expansion method. This work was supported by NIH grants R01DC011099 from the National Institute on Deafness and Other Communication Disorders (NIDCD), R21NS085772 from the National Institute of Neurological Disorders and Stroke (NINDS), and RF1MH120016 from the National Institutes of Mental Health (NIMH) to A.E.P.

## Author Contributions

All authors contributed to experimental design, each with particular emphases. S.P.C.G. performed and analyzed the expansion experiments reported here; S.I. performed initial expansion experiments and performed standard immunochemical experiments. A.E.P. oversaw all aspects of the project. All authors wrote the paper and approved of the manuscript.

## SUPPLEMENTAL FIGURE LEGENDS

**Supplemental Figure 1.**
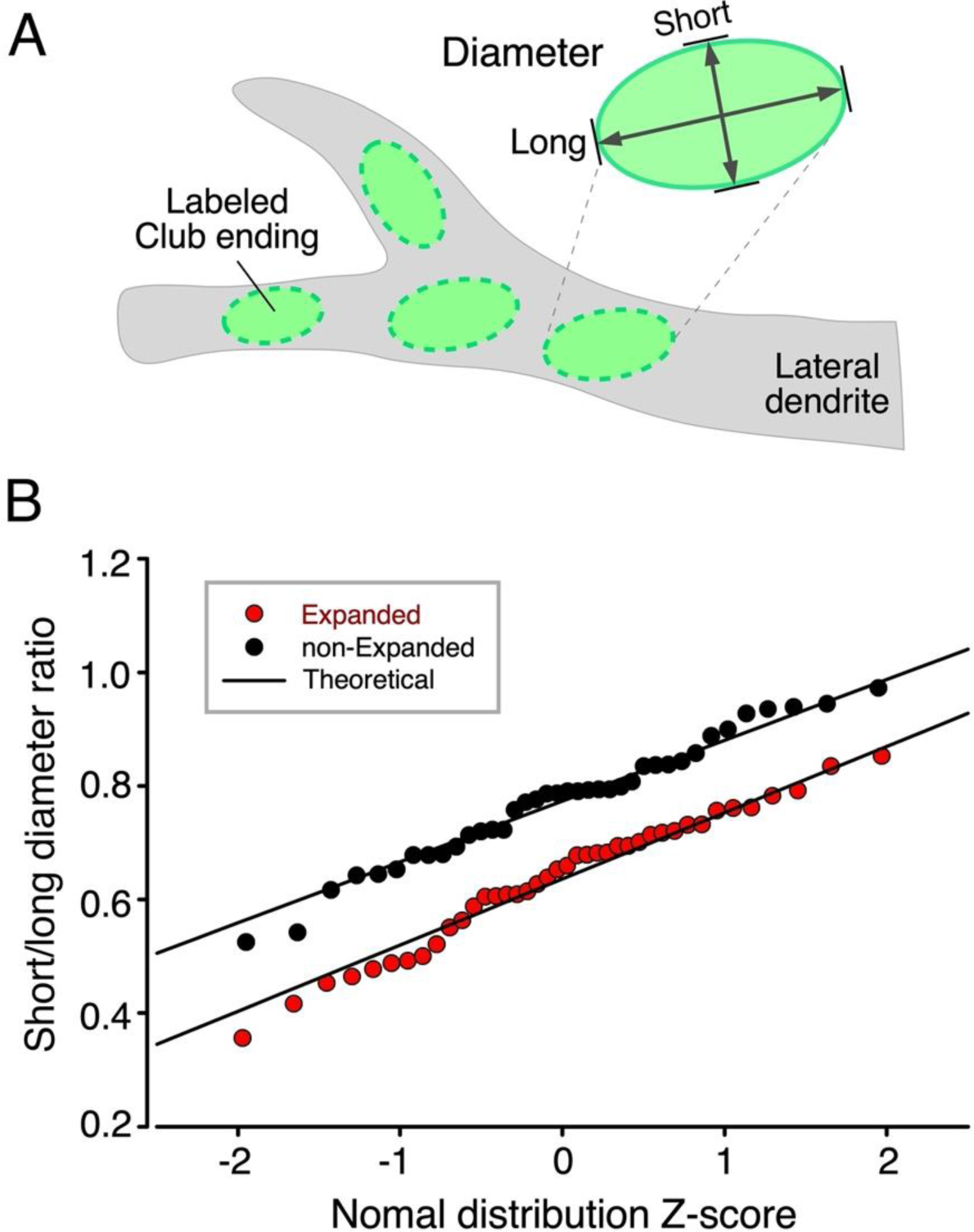
Short-long diameter ratio probability plots. (A) Cartoon illustrates labeled CE contact areas in the lateral dendrite of the M-cell and the measurement of their short and long diameters. (B) Short-long diameter ratios obtained from expanded (red circles) and non-expanded (black circles) CE contact areas are shown plotted against their standardized normal Z-scores calculated from sampled cumulative probabilities. Straight lines (Theoretical) show data expected from normally-distributed short-long ratios derived from sample means and standard deviations. In both expanded and non-expanded cases, experimental distributions fit theoretical normal distributions well. It may also be observed that the variability of ratios is nearly identical for expanded and non-expanded data, indicating that the expansion process had no selective effects across the CE population.

